# A Regularized Bayesian Dirichlet-multinomial Regression Model for Integrating Single-cell-level Omics and Patient-level Clinical Study Data

**DOI:** 10.1101/2024.06.04.597391

**Authors:** Yanghong Guo, Lei Yu, Lei Guo, Lin Xu, Qiwei Li

## Abstract

**Summary:** The abundance of various cell types can vary significantly among patients with varying phenotypes and even those with the same phenotype. Recent scientific advancements provide mounting evidence that other clinical variables, such as age, gender, and lifestyle habits, can also influence the abundance of certain cell types. However, current methods for integrating single-cell-level omics data with clinical variables are inadequate. In this study, we propose a regularized Bayesian Dirichlet-multinomial regression framework to investigate the relationship between single-cell RNA sequencing data and patient-level clinical data. Additionally, the model employs a novel hierarchical tree structure to identify such relationships at different cell-type levels. Our model successfully uncovers significant associations between specific cell types and clinical variables across three distinct diseases: pulmonary fibrosis, COVID-19, and non-small cell lung cancer. This integrative analysis provides biological insights and could potentially inform clinical interventions for various diseases.

## 1. Introduction

Single-cell RNA sequencing (scRNA-seq) has emerged as a powerful tool for discerning cell types within complex tissues and elucidating their functional roles (Kolodziejczyk et al., 2015; Papalexi and Satija, 2018; Luecken and Theis, 2019). However, translating cell type abundance into phenotypic associations is increasingly recognized as contingent upon multifaceted clinical variables such as age, gender, race, and ethnicity categories, among others. Recent scientific breakthroughs underscore the profound influence of these factors on modulating the abundance of specific cell types (Newman et al., 2019), yet existing methodologies for integrating scRNA-seq data with clinical variables are still inadequate. Therefore, there is an increasing need for innovative statistical approaches that can effectively integrate single-cell-level omics data with diverse clinical variables, thereby enhancing our understanding of the intricate relationships between cellular composition and phenotypic traits in scRNA-seq studies.

The integration of biological profiles (e.g., microarray data, bulk RNA-seq data, metagenomics data, etc) and clinical data has long been of interest. For instance, Gevaert et al. (2006) proposed a Bayesian network to integrate the microarray and clinical data for predicting the prognosis of breast cancer. Zhu et al. (2017) integrated clinical and multiple omics data for prognostic assessment across human cancers with the help of a kernel learning method. Li et al. (2017) developed a multivariate zero-inflated logistic normal model to quantify the associations between microbiome abundances and multiple factors based on microbiome compositional data instead of the count data. The analysis of sequence count data, particularly when integrating clinical variables, has been addressed through various well-established statistical methods. For instance, Wadsworth et al. (2017) introduced an integrative Bayesian Dirichlet-multinomial (DM) regression model tailored for microbiome sequencing reads. Jiang et al. (2021) developed a zero-inflated negative binomial (NB) regression model for similar datasets, utilizing the paired taxonomic tree structure to enhance the integrative analysis. The renowned DESeq2 package (Love et al., 2014) employs the NB distribution for analyzing bulk RNA-seq data. These models typically rely on Poisson or NB distributions for count data, with the DM distribution gaining popularity due to its effectiveness in characterizing the compositionality of some sequence count data (e.g., microbiome data). Zero-inflated distributions have been employed to address the prevalent issue of data sparsity, such as in microbiome and scRNA-seq datasets. However, the specific investigation of the association between clinical variables and scRNA-seq data, as well as its profound implications for certain diseases and biological significance, remains underexplored.

In this study, we employ a DM log-linear regression model to analyze cell type abundance based on scRNA-seq data alongside relevant clinical variables. Drawing upon the previous works (Wadsworth et al., 2017; Jiang et al., 2021), our approach facilitates the examination of associations between cell types and various clinical variables. We adopt spike-and-slab priors for the regression coefficients to selectively identify significant relationships. The efficacy of our model is demonstrated through a simulation study and further validated using three real datasets from distinct diseases. To better elucidate the connections between cell type abundance derived from scRNA-seq data and the related clinical variables, we construct a hierarchical tree approach that highlights the impact of these clinical variables on different levels of cell types. Exploring the association between clinical variables and cell types offers valuable insights into the role of cell abundance in defining phenotypes and the mechanisms through which these variables affect cellular dynamics. This understanding is crucial for deciphering the complex interplay between cellular composition and external factors, thereby advancing our knowledge of cellular function and its impact on broader biological systems.

The article is organized as follows. Section 2 introduces the data preprocessing steps that generate the cell-type abundance data and outlines the data notations. Section 3 describes the structure of the Bayesian DM log-linear regression model, incorporating spike-and-slab priors to enhance model robustness. Section 4 details the Markov chain Monte Carlo (MCMC) algorithm and the posterior inference of the key model parameters. In Section 5, we evaluate the model’s performance through a simulation study and present the results from three case studies involving pulmonary fibrosis, COVID-19, and non-small cell lung cancer. Our conclusions are presented in Section 6.

## 2. Data

### 2.1 Data preparation

The raw scRNA-seq data, consisting of short reads from the transcript 3’ end with unique molecular identifiers (UMIs) for each cell type, are processed using the Cell Ranger Pipeline (v6.1.1) for sample demultiplexing, barcode processing, and single-cell gene counting matrix generation. Raw reads of FASTQ files are aligned to the human hg38 reference genome. The Seurat package (v4.1.1)(Hao et al., 2021) is used for clustering and uniform manifold approximation and projection (UMAP) analysis. Differentially expressed genes (DEGs) are found using the Wilcoxon rank-sum test with a *p*-value threshold of ⩽ 0.05, and *p*-values are adjusted based on Bonferroni correction for multiple comparisons. Cell-type specific marker genes are identified for cell type assignment with the FindConservedMarkers function in the Seurat package, and then using the DESeq2 function to identify DEGs. Data are then normalized and scaled to account for technical variations in the Seurat package. Next, we perform dimension reduction with the UMAP algorithm to visualize the data in a lower-dimension space. The clustering algorithm Louvain is applied to group cells with similar expression profiles into clusters. Each cluster is examined for DEGs, known as marker genes, which are characteristic of specific cell types. By comparing these marker genes to known gene expression profiles from established cell type databases or literature, we are able to assign each cluster to a specific cell type.

### 2.2 Data notation

After the cell-type information has been obtained from the above steps, we generate an *N* × *P* cell-type abundance count matrix ***Y***, where each row indexed by *i* (*i* = 1, …, *N*) represents a patient and each column indexed by *j* (*j* = 1, …, *P*) corresponds to a specific cell type. Each row ***y***_*i*_ = [*y*_*i*1_, …, *y*_*iP*_] indicates the cell-type abundance of patient *i*, with *y*_*ij*_ being the total number of cell type *j* found in the scRNA-seq data of patient *i*. Besides, we summarize the paired clinical data as an *N* × *R* matrix denoted by ***X***, with the *i*-th row ***x***_*i*_ = [*x*_*i*1_, …, *x*_*iR*_] representing the measurements of all the *R* clinical variables from patient *i*.

## 3. Model

To identify significant associations between a range of clinical variables and cell types or grouped cell types, we introduce a hierarchical Bayesian framework that combines the analysis of cell-type abundance count data with clinical information. In this framework, cell type abundance from a patient is assumed to be drawn from a DM distribution. Clinical variables are seamlessly incorporated into the model by parameterizing the DM distribution’s parameters through a log-linear regression approach. This methodology enables a direct and integrated examination of how clinical variables influence cell-type distributions.

### 3.1 Dirichlet-multinomial (DM) level

We start by modeling each row of the cell-type abundance data ***Y*** with a multinomial distribution

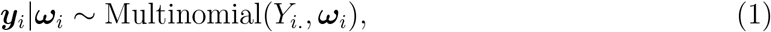

with 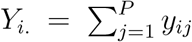 being the summation of all cell-type counts in vector ***y***_*i*_, and the *P*-dimensional vector ***ω***_*i*_ = [*ω*_*i*1_, …, *ω*_*iP*_]^⊤^ is defined on a *P*-dimensional simplex

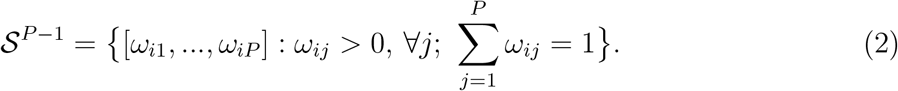

We further impose a conjugate Dirichlet prior on the parameter ***ω***_*i*_ to allow for over-dispersed distributions, that is ***ω***_*i*_|***α***_*i*_ ~ Dirichlet(***α***_*i*_), where each element of the *P*-dimensional vector ***α***_*i*_ = [*α*_*i*1_, …, *α*_*iP*_]^⊤^ is strictly positive. By integrating ***ω***_*i*_ out, we get the resulting DM model, ***y***_*i*_|***α***_*i*_ ~ DM(***α***_*i*_), where the corresponding probability mass function is

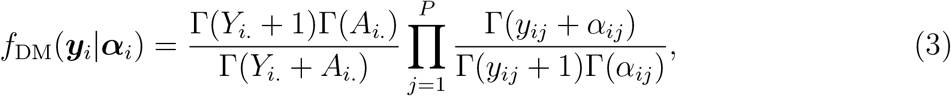

where 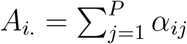. Compared with multinomial distribution, this setting allows for over-dispersed distributions by inducing an increase in the variance by a factor of (*Y*_*i*._+*A*_*i*._)/(1+*A*_*i*._), which is greater than 1.

### 3.2 Log-linear regression level

The covariates matrix ***X*** are then incorporated into the model by a log-linear regression framework, where the parameter ***α***_*i*_ of the DM distribution is linked to the covariates by specifying

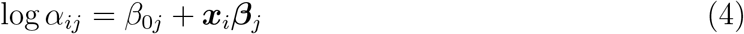

where ***β***_*j*_ = [*β*_1*j*_, …, *β*_*Rj*_]^⊤^ is an *R*-dimensional vector, with each element *β*_*rj*_, *r* = 1, …, *R* modeling the effect of the *r*-th covariate on the *j*-th cell type. The intercept term *β* _0*j*_ serves as the log baseline parameter for the cell type *j*.

Identifying the significant associations between cell type abundance and clinical variables is equivalent to finding t he n on-zero *β*_*rj*_. I n p ractice, n ot a ll o f t he c linical variables are associated with the abundance of each cell type, therefore, we specify a spike-and-slab prior as

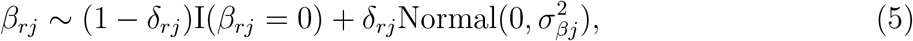

where δ_*rj*_ = 1 indicates the *r*-th covariate is associated with the abundance of the *j*-th cell type, and δ_*rj*_ = 0 otherwise. Here I(·) is an indicator function. The latent binary variables δ_*rj*_ serve as the indicators of which pairs of cell type and clinical variable have a significant association. We further complete the model by setting the prior of 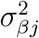 to be an inverse-gamma (IG) distribution IG(*a*_*β*_, *b*_*β*_). A common way of choosing the value of *a*_*β*_ and *b*_*β*_ is to set *a*_*β*_ = 2 and *b*_*β*_ = 10, which suggests a flat prior of variance 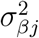 that encourages the selection of relatively large effects.

The model is completed by setting *β*_0*j*_ ~ Normal 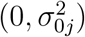 and δ_*rj*_ ~ Beta-Bernoulli(*a*_*p*_, *b*_*p*_). Practically, we set *a*_*p*_ + *b*_*p*_ = 2, and the choice of hyperparameters *a*_*p*_ and *b*_*p*_ reflects the prior belief that a proportion *a*_*p*_*/*(*a*_*p*_ + *b*_*p*_) of the cell type and covariate associations would be selected as discriminating among all pairs. For most cases, a value of *a*_*p*_*/*(*a*_*p*_ + *b*_*p*_) ∈ [0.1, 0.2] corresponds to assuming a priori that 10% to 20% of the covariates will be selected. Finally, we further complete the model by setting 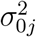 as a relatively large value (e.g., 10), since such a choice suggests a flat prior distribution on the location of the coefficients. We denote **Δ** = [δ_*rj*_]_*R*×*P*_ and ***B*** = [*β*_*rj*_]_*R*×*P*_, where *r* = 1, …, *R, j* = 1, …, *P*, and ***β***_0_ = [*β*_01_, …, *β*_0*P*_]^⊤^.

### 3.3 Incorporating cell-type tree structure

Cell-type abundance data can be summarized at different hierarchical levels based on their similarities. Given all the cell types (leave nodes) at the bottom level, we can build a binary tree from bottom to top by the “height” and “merge” attributes of each parent node (Figure 1a). For example, leaf (cell type) *j* and *j* + 1 merge first with the smallest height, then we obtain the first parent node, denoted by “node 1”, and the corresponding “layer 1”. *P* is the total number of cell types as in the count matrix ***Y***. From now on, “node” only refers to the parent nodes, not the leaves.

**Figure 1:**
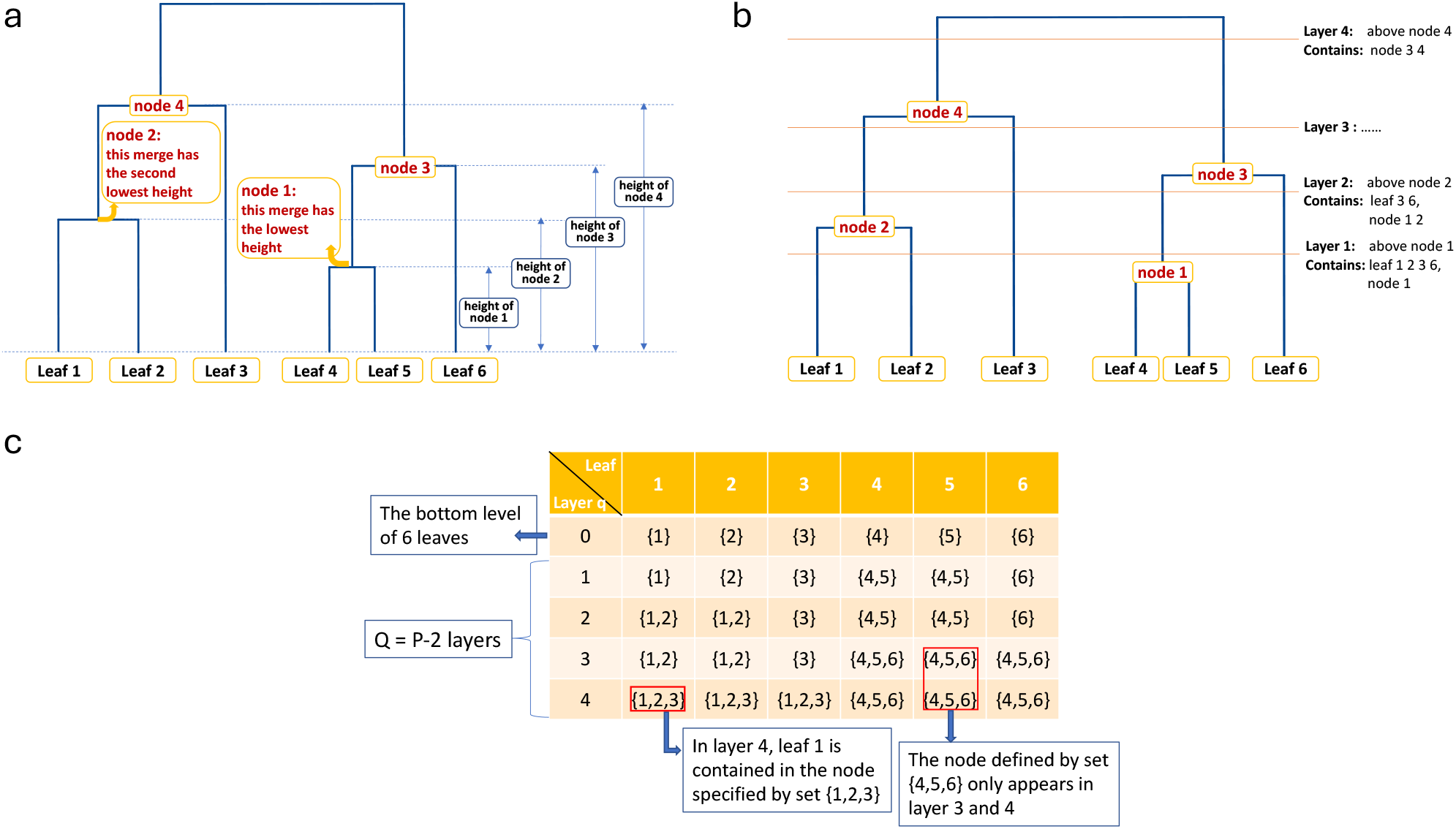
Tree structure explanation. (a) The parent nodes are ranked by their height. (b) The tree is cut into layers. (c) An example of matrix ***M***_(*Q*+1)*×P*_.

By merging the nodes in this manner, we end up having *Q* = *P* − 2 parent nodes from bottom to top. By cutting the tree horizontally from the bottom, for each merge we get a new cut that represents a new layer, and we have in total *Q* layers (Figure 1b).

Now consider a (*Q* + 1) *× P* matrix ***M*** where each row ***m***_*q*_ = [𝒥_*q*1_, …, 𝒥_*qP*_] is a vector of sets representing the composition of nodes in layer *q, q* = 0, …, *Q. q* = 0 indicates the bottom level that contains only the leaves. 𝒥_*qj*_ ⊂ {1, …, *P*}, *j* = 1, …, *P* is a set of integers representing all the leaves in a certain node. Taking Figure 1c as an example, for layer *q* = 3, leaf 1 and 2 are in the same parent node denoted by 𝒥_31_ = {1, 2}, and leaf 4, 5, and 6 are in the node denoted by 𝒥_34_ = {4, 5, 6}.

We can aggregate the counts ***Y*** into any upper layer *q* based on the way the nodes merge, denoted by an *N ×* (*P* − *q*) matrix ***Y*** ^(*q*)^. Notice that ***Y*** ^(0)^ = ***Y***. More explicitly, let *θ*(·) be a function of vectors that can extract all the unique values in the input vector, then *θ*(***m***_*q*_) returns all the unique sets in ***m***_*q*_, and by Figure 1a and 1c we can observe that there should be *P* − *q* unique sets in *θ*(***m***_*q*_). Then ***Y*** ^(*q*)^ is obtained in the following manner: for row ***y***^(*q*)^ in ***Y*** ^(*q*)^, each element of it is calculated by

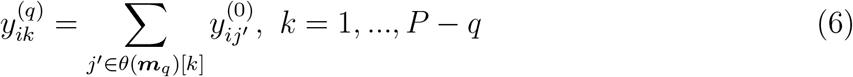

where *θ*(***m***)[*k*] is the *k*th unique set in *θ*(***m***), and 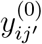 is the count in row *i* and column *j*^*′*^ of ***Y*** ^(0)^. In Figure 1c, for layer 3 we have *θ*(***m***_3_) = {{1, 2}, {3}, {4, 5, 6}}.

Correspondingly, assume 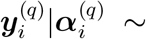 Dirichlet 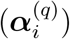, we link the parameter with the covariates by

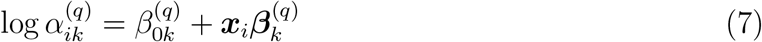

where 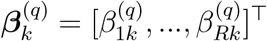 is an *R*-dimensional vector, with each element 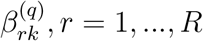 modeling the effect of the *r*-th covariate on the node specified by set *θ*(***m***_*q*_)[*k*]. Similarly, we specify a spike-and-slab prior as 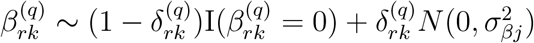.

For a specific node *u* defined by set 𝒥_*u*_ ⊂ {1, …, *P*}, the corresponding matrix of indicators and regression coefficients are calculated as

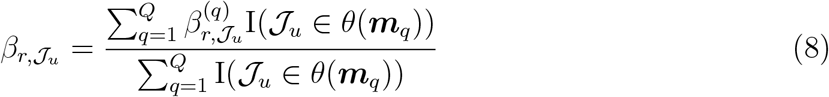

and

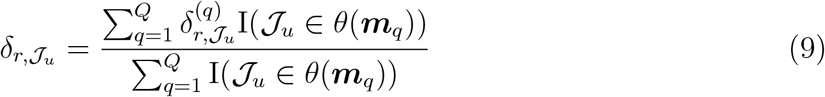

where 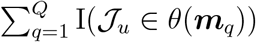 indicates the total number of appearance of node *u* in all *Q* layers, and 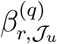 is the regression coefficient for layer *q* that is associated with the *r*th covariate and node *u*.

## 4. Model Fitting

### 4.1 The MCMC algorithm

We implement the MCMC algorithm for posterior inference by updating each step with Random Walk Metropolis-Hasting (RWMH) sampling. The details of the MCMC algorithm are in Algorithm 1.

#### Algorithm 1

The details of the MCMC algorithm

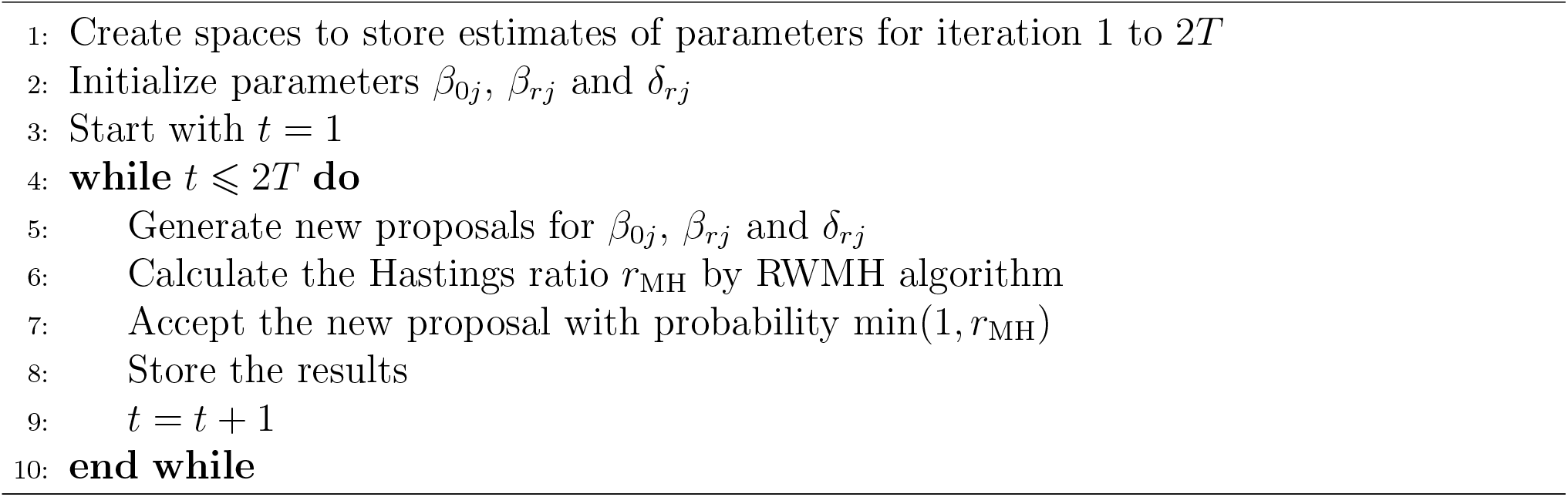

According to the model described in Section 3, the full data likelihood is given as follows.

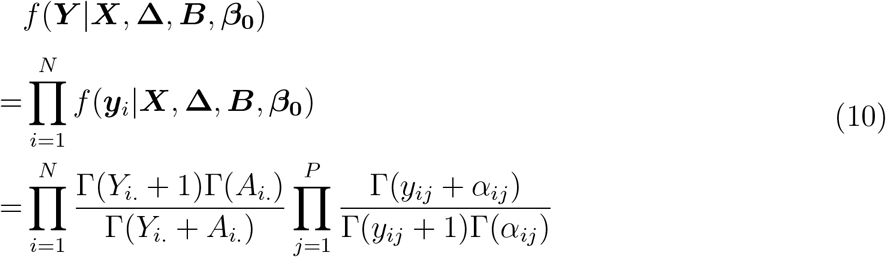

where 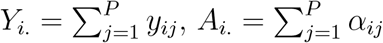, and log *α*_*ij*_ = *β*_0*j*_ + ***x***_*i*_***β***_*j*_.

Then, we update the parameters in each iteration following the steps below:

**Jointly update** δ_*rj*_ **and** *β*_*rj*_ We perform a between-model step first using an *add-delete* algorithm. For each *j* = 1, …, *P* and each *r* = 1, …, *R*, we change the value of δ_*rj*_. For the *add* case, i.e. δ_*rj*_ = 0 → δ_*rj*_ = 1, we propose 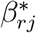 from 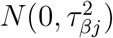. For the *delete* case, i.e. δ_*rj*_ = 1 → δ_*rj*_ = 0, we set 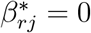. Matrices **Δ**^∗^ and ***B***^∗^ are identical to **Δ** and ***B***, except for the elements δ_*rj*_ and *β*_*rj*_, which are replaced by 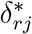 and 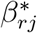, respectively. We then accept the proposed new 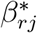 with probability min(1, *r*_MH_), where

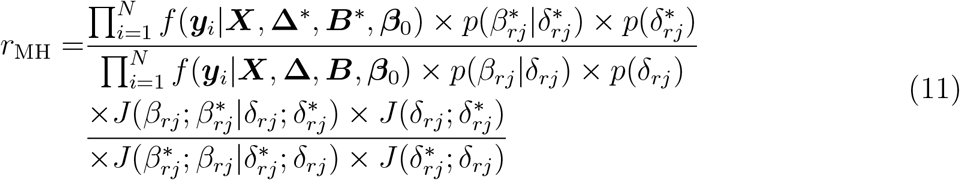

**Update** *β*_*rj*_ **when** δ_*rj*_ = 1: A within-model step is followed to further update each *β*_*rj*_ where δ_*rj*_ = 1. We first propose a new 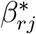 from 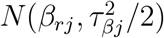 with RWMH algorithm. ***B***^∗^ is identical to ***B***, except that the element *β*_*rj*_ is replaced by 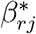. Then we accept the proposed value with probability min(1, *r*_MH_), where

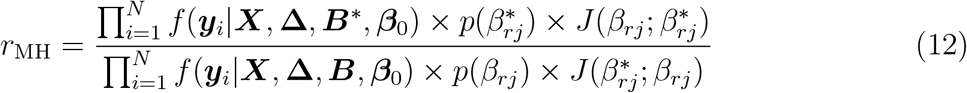

**Update** *β*_0*j*_: We update each *β*_0*j*_, *j* = 1, …, *P* sequentially using RWMH algorithm. We first propose a new 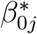 from 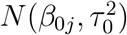. 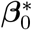 is identical to ***β***_0_, except that the element *β*_0*j*_ is replaced by 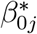. Then we accept the proposed value with probability min(1, *r*_MH_), where

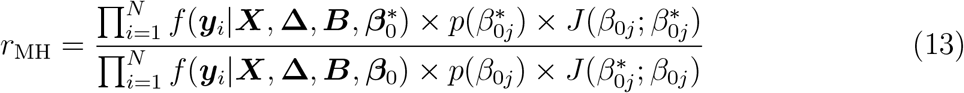

### 4.2 Posterior inference

For the posterior inference, our aim is to identify the significant associations between cell type abundance and covariates by selecting over **Δ** and the corresponding matrix ***B***. One way to summarize the posterior inference of the latent variable **Δ** is via the estimates of the marginal posterior probabilities of inclusion (PPI). Suppose that we have in total 2*T* iterations and the burn-in rate is 50%, then the PPI of each single δ_*rj*_ is calculated by 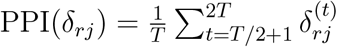, where 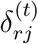 is the accepted proposal of δ_*rj*_ in the *t*th iteration. In this way, we are able to select the significant associations by specifying a threshold on PPIs. One choice of threshold is to assign it a fixed value, e.g., 0.5. Another more popular way is to choose a threshold that controls the Bayesian false discovery rate (FDR) *γ* which is calculated as

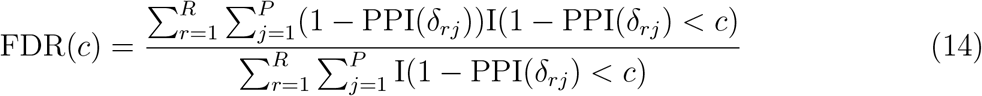

where *c* is the threshold and I(·) is the indicator function. An optimal choice of *c* can be found for a certain error rate *γ* by choosing *c*^*′*^ such that FDR(*c*^*′*^) *< γ*, and a common setting of the error rate is *γ* = 0.05.

## 5. Results

To illustrate the capability of our model to estimate associations between cell type abundance and clinical variables, we applied it to a simulated dataset. We conducted comparative analyses with established classical methods. The superior performance of our model underscored its efficacy in addressing integration challenges. Then we applied the model to three real datasets. Compared to the results in the original studies, our model effectively highlighted relationships between cell types and variables. Additionally, it shed light on new findings not identified by the original studies, further indicating our model’s effectiveness.

### 5.1 Simulation study

We first evaluated the proposed model with a simulated dataset. To mimic the real-world scenario, we utilized the covariate matrix ***X*** from the first real dataset of our study (the pulmonary fibrosis dataset). In this dataset, *N* = 25 and *R* = 4, which indicates that there are 25 patients and four covariates under consideration. We set the number of cell types to be *P* = 30 and simulated the ***Y*** in the following manner. We first sampled all the entries of matrix **Δ** from Bernoulli(0.2). When the corresponding δ_*rj*_ = 0, *β*_*rj*_ was set to zero, and when the corresponding δ_*rj*_ = 1, we sampled *β*_*rj*_ by first sampling a 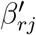 from a uniform distribution Uniform(1, 5), and a corresponding parameter *z* from Bernoulli(0.5). If *z* = 1, 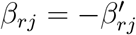, and if 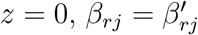, which means *β*_*rj*_ has a 50% probability of being negative.

The vector ***β***_0_ was sample from a truncated normal distribution Normal(0, 5) within the limit [−3, 3]. Each row of ***Y***, denoted by ***y***_*i*_, was simulated from Multinomial(*Y*_*i*._, ***ω***_*i*_), where ***ω***_*i*_ ~ Dirichlet(***α***_*i*_). ***α***_*i*_ can be calculated by log *α*_*ij*_ = *β*_0*j*_ + ***x***_*i*_***β***_*j*_. *Y*_*i*._ ~ Uniform(300, 7, 000), representing the total number of cells from each patient.

We implemented the proposed Bayesian model by setting *a*_*p*_ = 0.4, *b*_*p*_ = 1.6, which corresponds to assuming a priori that 20% of the covariates would be selected. We set *a*_*β*_ = 2, *b*_*β*_ = 10, which suggests a flat prior distribution of *β*_*rj*_ that encourages the selection of relatively large effects. We further set 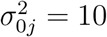 to allow a flat prior on *β*_0*j*_. The variance of the proposal distribution of the MCMC algorithm was set as 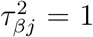. We set the total iteration of MCMC as 2*T* = 20, 000, and the burn-in rate as 50%. The receiving operating curves (ROC) plot and the corresponding area under the curve (AUC) between the true values and the PPI of **Δ** served as the metric of this study.

Then we compared our model to two simple alternatives. First, we fit a simple linear regression model on each column ***y***_*j*_ of ***Y*** (each column represents the counts of a single cell type among patients), using ***X*** as the covariate matrix. We then obtained *P* sets of regression coefficients. Subsequently, we constructed the ROC curve and calculated the AUC using the *p*-values associated with each regression coefficient. Next, we tested the FDR-corrected correlation coefficient of each pair of columns between ***Y*** and ***X*** (i.e. pair-wise correlation tests) and saved the adjusted *p*-values for the calculation of ROC and AUC. The comparison results are shown in Figure 2. Our model reached an AUC of 0.995, while the correlation test and linear regression obtained AUC of 0.832 and 0.808, respectively.

**Figure 2:**
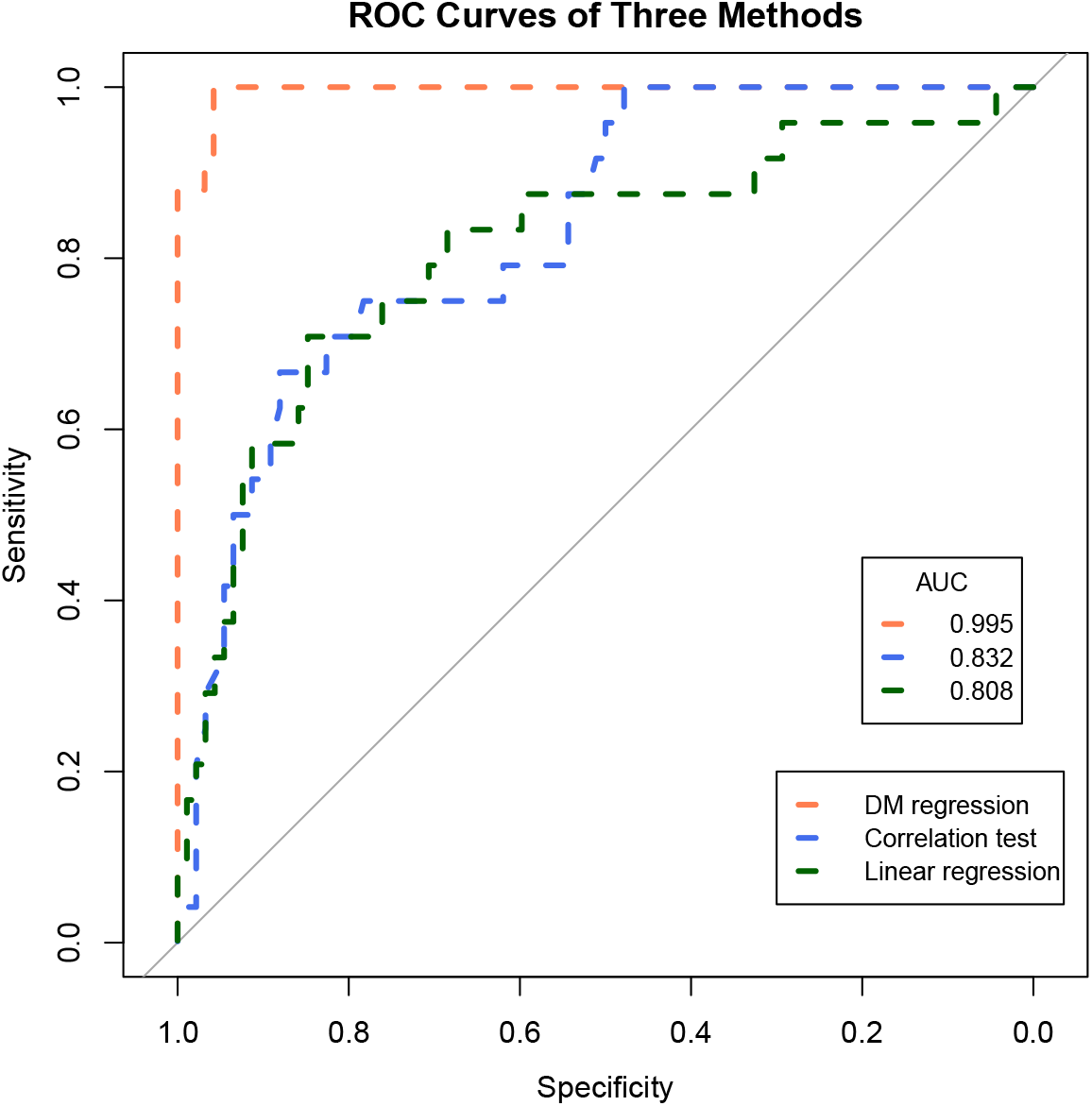
Simulation study result. The ROC and AUC results on the simulated dataset with three methods: DM log-linear regression, pair-wise correlation test, and simple linear regression.

### 5.2 Application to pulmonary fibrosis dataset

Pulmonary fibrosis (PF) is a chronic lung condition characterized by aberrant epithelial restructuring and the accumulation of extracellular matrix (ECM), marking a significant pathology within the pulmonary system. We applied our method to a scRNA-seq dataset from 30 patients, among whom 20 were diagnosed with PF, while the remaining ten, who didn’t have PF, served as the control group (Habermann et al., 2020). The authors have previously defined *P* = 30 cell types in this PF scRNA-seq data and also provided clinical information on these 30 patients, including age, sex, smoking history, and disease status (*R* = 4). The patients were between ages of 17 and 74. Among these 30 patients, 17 (56.7%) were male, and 16 (53.3%) had ever smoked in their lives. After removing the patients with missing values, we kept *N* = 25 patients to proceed with our study.

Figure 3a depicts the associations between the abundance of 30 distinct cell types and four clinical parameters, namely age, sex, smoking history, and disease status (i.e., with PF or not). The analysis revealed positive correlations between certain cell types [e.g., Epi(KRT5-/KRT17+) and Epi(basal)] and the presence of PF. Conversely, we observed no significant correlations between these cell types and other clinical factors, such as age, sex, or smoking history. Figure 3b presents the posterior estimates of the regression coefficients *β*_*rj*_ between disease status and the aforementioned cell types. The significant coefficients corroborate the associations highlighted in Figure 3a, emphasizing the relationship between changes in disease status and alterations in cell type abundances. Figure 3c presents the 95% posterior credible intervals for *β*_*rj*_ between the disease status and the cell types Epi(KRT5-/KRT17+) and Epi(basal), respectively. These exclusively positive intervals further substantiate their strong positive association with PF. Figure 3d visualizes the 95% posterior credible intervals for significant regression coefficients 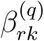 within the hierarchical tree structure. These intervals underscore the significance of these coefficients in elucidating the impact of PF on cell type distributions. Figure 3e highlights elevated levels of Epi(SCGB1A1+), Epi(SCGB3A2+), and Epi(basal) cell types in patients diagnosed with PF. This suggests a significant increase in the abundance of these cell types among individuals with the disease, as discerned through the hierarchical model. Finally, Figure 3f displays the predicted proportions of Epi(basal) and Epi(KRT5-/KRT17+) alongside actual proportions in patients. The close alignment between the predicted and actual proportions underscores the model’s accuracy and emphasizes the heightened presence of these cell types in PF patients.

**Figure 3:**
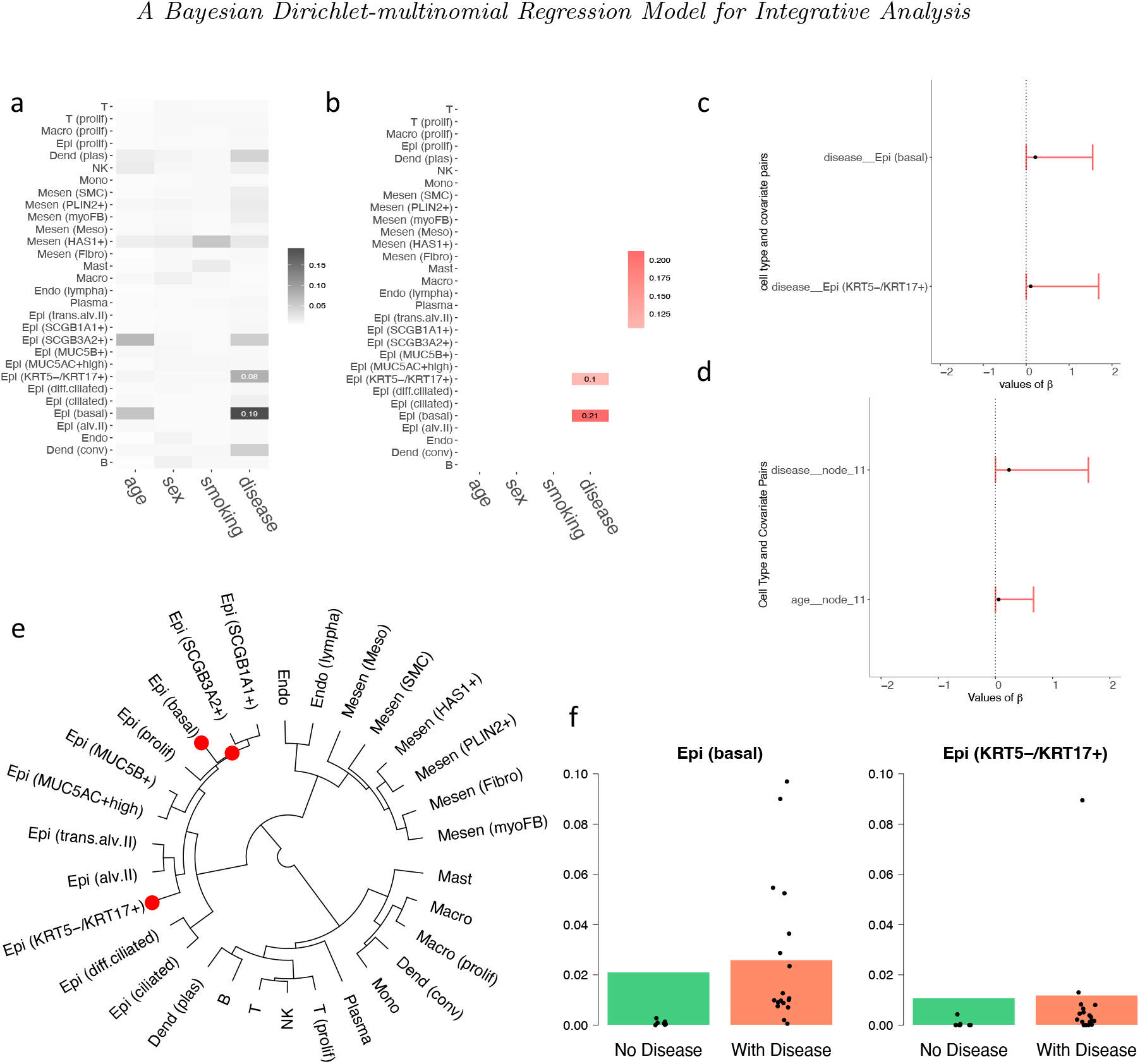
Results on the PF dataset. (a) Heatmap showing the relationship between covariates and cell types, with darker colors indicating stronger relationships. (b) Heatmap of estimated regression coefficients, where red indicates a positive correlation. (c) 95% credible interval for the significant *β*_*rj*_ chosen by the threshold controlling the FDR *<* 0.05. (d) 95% credible interval for the significant 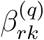 of the tree structure chosen by the threshold controlling the FDR *<* 0.05. (e) Cladograms of the identified cell types, where red dots represent the nodes of cell types that have a significantly higher abundance when diagnosed with PF. (f) The underlying barplot represents the predicted proportions (*α*) of the certain cell type by the model, with black dots showing the actual proportions of this cell type in each patient in the dataset.

It is worth noting that our predicted associations are consistent with multiple known biological observations: Basal cell hyperplasia is indeed a characteristic feature observed in epithelial remodeling associated with fibrotic lung disease and other chronic lung conditions. This phenomenon involves an increase in the number of basal cells in the epithelium, which is the layer of cells lining the respiratory tract (Beppu et al., 2023; Ortiz-Zapater et al., 2022). The aberrant expansion of KRT5-/KRT17+ epithelial cells in pulmonary fibrosis lungs is important in understanding the pathogenesis of fibrotic lung diseases. This observation highlights changes in the epithelial cell populations that contribute to the progression of these diseases (Habermann et al., 2020; Valenzi et al., 2021). Previous papers have validated the positive correlation between Epi (basal) and Epi (KRT5-/KRT17+) abundance with pulmonary fibrosis, supporting the power of DM regression in revealing the genuine biological relationship between cell type abundance and disease.

### 5.3 Application to COVID-19 dataset

Immune dysregulation in patients with coronavirus disease 2019 (COVID-19) significantly influences the symptom and mortality rates. To provide a comprehensive landscape of relevant immune cell dynamics in COVID-19 patients, a recent study leveraged scRNA-seq to analyze 284 samples from 196 individuals, including COVID-19 patients and controls, thereby elucidating a comprehensive immune landscape that comprises 1.46 million cells across *P* = 12 distinct immune cell types: B, CD4, CD8, dendritic cells (DC), epithelial cell (Epi), macrophages (Macro), mast cell (Mast), megakaryocytes (Mega), monocytes (Mono), neutrophils (Neu), natural killer cells (NK), and plasma (Ren et al., 2021). We analyzed five clinical variable: age, sex, scRNA-seq platform, disease symptoms, and disease stages. By combining the disease symptom and stage into a single variable, we run our model with *R* = 4 clinical variables. After preprocessing the data by removing samples with missing values, *N* = 283 samples were kept in our study, and 177 (62.5%) were from male patients. The ages range from 6 to 92, and two different sequencing platforms, 10 *×* 3^*′*^ and 10 *×* 5^*′*^, were utilized to generate the scRNA-seq data. 28 (9.9%) samples served as the control group, 121 (42.8%) exhibited mild to moderate symptoms, and 134 (47.3%) had severe symptoms. Additionally, 139 (49.1%) samples were collected from patients at the convalescence stage, and 116 (41.0%) during the progression stage.

As shown in Figure 4a, among these immune cell types, NK, Mono, DC, CD8, and CD4 were found to have a higher abundance level within patients exhibiting disease progression and severe symptoms. Notably, Figures 4b and 4c underscore a significant observation: age displays a negative correlation with the abundance of CD4 and CD8 T cells, particularly evident in patients undergoing disease progression and exhibiting moderate symptoms. Figure 4b (left) delineates a significant increase in the cell types comprising Mast, Macro, DC, and Mono among patients in advancing disease stages. Conversely, Figure 4b (right) demonstrates that all hierarchical cell types are more abundant in patients with severe symptoms, highlighting the impact of disease severity on immune cell dynamics. Further elaborating on these associations, Figure 4c confirms the inverse relationship between age and the levels of CD4 and CD8 expression, aligning with the observed heightened response in critical conditions. During severe disease progression, increased levels of NK, Mono, DC, and both CD8 and CD4 T cells are evident, indicating an intensified immune response. Figure 4d illustrates the 95% posterior credible intervals for the significant regression coefficients (*β*_*rj*_) between the disease status, symptom severity, and the aforementioned immune cells. The intervals substantially exceeding zero for CD4, CD8, Mono, and NK suggest a robust positive association with advanced disease states, highlighting their pivotal roles in the immune system’s response to disease progression. Moreover, Figure 4e presents the 95% posterior credible intervals for the significant 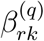 of the hierarchical tree structure, further delineating the statistical significance of these relationships within the clustered groups of immune cells. Finally, Figure 4f shows the predicted proportions of CD4, CD8, Mono, and NK under various clinical scenarios, with the actual proportions observed in patients overlaid as black dots. This visualization not only confirms the model’s accuracy but also emphasizes the prevalence of these immune cells in patients undergoing severe disease progression.

**Figure 4:**
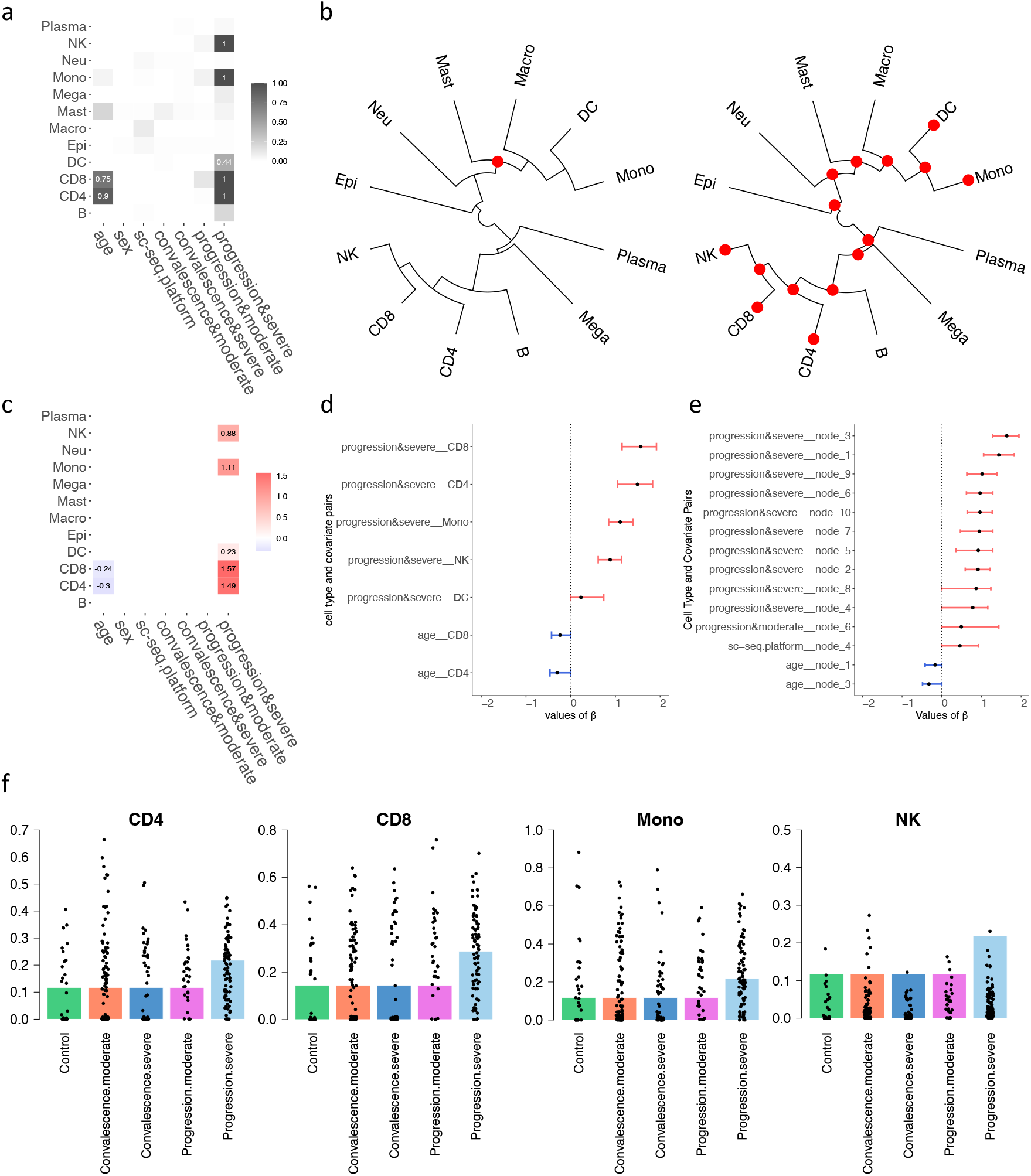
Results on the Covid-19 dataset. (a) Heatmap showing the relationship between covariates and cell types, with darker colors indicating stronger relationships. (b) Cladogram of the identified cell types, sampled during the progression stage and categorized by severity (left for moderate and right for severe). Red dots indicate nodes of cell types with significantly higher abundance, while blue represents lower abundance. (c) Heatmap of estimated regression coefficients, where red indicates a positive correlation, and blue denotes a negative correlation. (d) 95% credible interval for the significant *β*_*rj*_ chosen by the threshold controlling the FDR *<* 0.05. (e) 95% credible interval for the significant 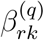 of the tree structure chosen by the threshold controlling the FDR *<* 0.05. (f) The underlying barplot represents the predicted proportions (*α*) of the certain cell type by the model, with black dots showing the actual proportions of this cell type in each sample in the dataset.

Furthermore, it is noteworthy that our model predictions are consistent with biological facts: age-related thymic involution and the accumulation of memory T cells contribute to the decline of CD4+ and CD8+ T cells, which are essential components of the adaptive immune response. This phenomenon is termed immunosenescence and is associated with decreased immune function in older adults (Zhang et al., 2021; Ramasubramanian et al., 2022; Li et al., 2019). The negative correlation between CD8 and CD4 cell abundance with age is consistent with the biological truth.

### 5.4 Application to lung cancer dataset

Non-small cell lung cancer (NSCLC) is well-known for being a highly aggressive and heterogeneous disease with diverse histological subtypes. A comprehensive understanding of the immune and stroma cell types across various NSCLC patient subgroups with distinct phenotypes is still largely lacking. A recent study has applied scRNA-seq to delineate *P* = 33 distinct cell types sourced from *N* = 179 patients (Salcher et al., 2022). In our study, *R* = 4 covariates were examined, including age, sex, smoking history, and tumor stage. Of the 179 patients, 83 (46.4%) were male and 96 (53.6%) were female. Additionally, 101 (56.4%) patients had a history of smoking. Out of the 179 patients, 66 (36.9%) did not have NSCLC and served as the control group, 38 (21.2%) were in an advanced tumor stage, and 75 (41.9%) were in the early tumor stage.

The analysis depicted in Figures 5a and 5c reveals distinct patterns of cellular abundance that correlate with tumor progression and lifestyle factors. Specifically, patients in the early stages of tumor development exhibit increased levels of regulatory T cells (T(reg)), cytotoxic T cells (T(CD8+)), and helper T cells (T(CD4+)), alongside decreased abundances of monocytes (Mono) and alveolar macrophages (Macro(alv)). Conversely, patients with advanced tumors show reduced levels of Mono and Macro(alv), but an elevated presence of neutrophils. Additionally, a notable increase in plasmacytoid dendritic cells (Den(plas)), natural killer cells (NK), and B cells is observed among patients with a history of smoking, highlighting the impact of lifestyle factors on immune cell dynamics. Our hierarchical analysis, as demonstrated in Figure 5b, shows that the cell types comprising non-specific monocytes (Mono(non)), alveolar macrophages (Macro(alv)), macrophages (Macro), and dendritic cells expressing CD1c (Den(CD1c+)) are less abundant at both early and advanced stages of the disease. In contrast, early-stage tumors are characterized by a higher prevalence of T(reg), T(CD4+), T(CD8+), NK, Mast, Den(plas), B, conventional dendritic cells (Den(conv)), and Plasma, suggesting a more active immune response at this stage of tumor development. Figure 5d illustrates the 95% posterior credible intervals for significant regression coefficients (*β*_*rj*_) between disease covariates and cell types, establishing strong statistical evidence of their associations. Figure 5e further extends these findings by showing the credible intervals for significant 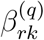 values within the hierarchical tree structure, providing insights into the interconnected nature of cell type dynamics and disease characteristics. In Figure 5f, we compared the estimated proportions with the observed proportions of these cell types, depicted through underlying barplots and overlaid black spots, respectively. This comparison reveals that Macro(alv) and Mono(non) are predominantly observed in non-cancer patients, whereas T cells, specifically T(CD4+), T(CD8+), and T(reg), are significantly more abundant in patients with early-stage lung cancer, underscoring their potential roles in early immune surveillance and response to tumor presence.

**Figure 5:**
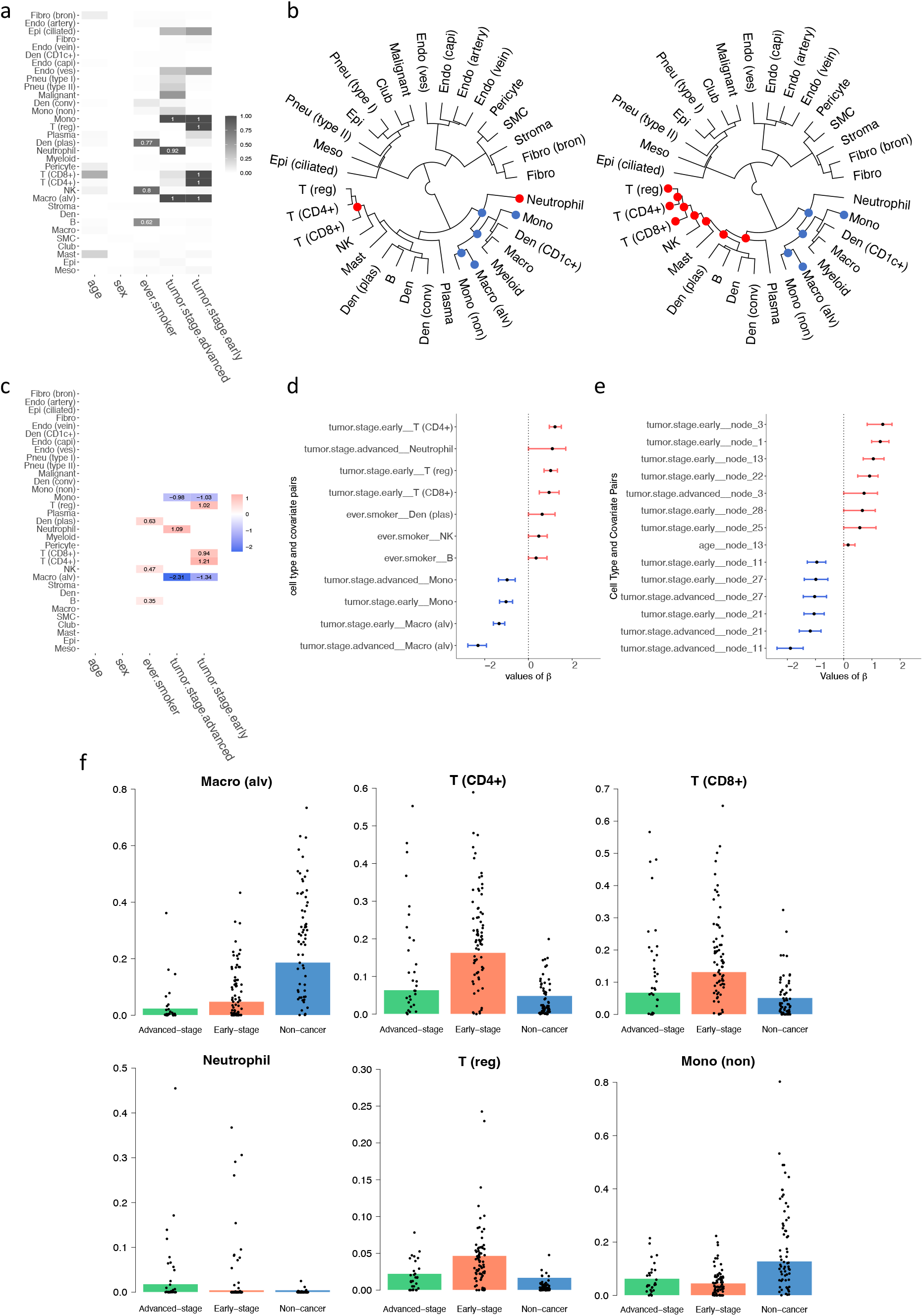
Results on the lung cancer dataset. (a) Heatmap showing the relationship between covariates and cell types, with darker colors indicating stronger relationships. (b) Cladogram of the identified cell types, with tumor stages shown as advanced (left) and early (right); red dots indicate nodes of cell types with significantly higher abundance, and blue signifies lower abundance. (c) Heatmap of estimated regression coefficients, where red indicates a positive correlation, and blue denotes a negative correlation. (d) 95% credible interval for the significant *β*_*rj*_ chosen by the threshold controlling the FDR *<* 0.05. (e) 95% credible interval for the significant 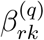 of the tree structure chosen by the threshold controlling the FDR *<* 0.05. (f) The underlying barplot represents the predicted proportions (*α*) of the certain cell type by the model, with black dots showing the actual proportions of this cell type in each patient in the dataset.

Furthermore, the predicted associations are consistent with published biological observations: T cells (T(reg)), cytotoxic T cells (T(CD8+)), and helper T cells (T(CD4+)) are crucial for anti-tumor immunity. In the early stage of lung cancer, regulatory T cells (Tregs) exhibit significant plasticity and functional diversity across various tumors within the tumor microenvironment and have been found to increase (Principe et al., 2021; Liang et al., 2022; Li et al., 2024). Several studies have demonstrated that tumors with a high quantity of FoxP3+ regulatory T cells (Tregs) also exhibit a substantial presence of other immune cells, including CD8+ cytotoxic T cells and proliferative immune cells (Koll et al., 2023; Oshi et al., 2022). Additionally, the increased count of CD4+ cells was observed in non-small cell lung cancer tumor-infiltrating lymphocytes (Woo et al., 2001). Studies reported that as lung cancer progresses, the proportion of alveolar macrophages (Macro(alv)) gradually decreases (Wang et al., 2023), which supports the negative coefficient of the abundance of Macro (alv) in both NSCLC early stage and advanced stage. Neutrophil expansion is associated with changes in the inflammatory milieu of patients with non-small cell lung cancer (NSCLC) who have resectable disease (Mitchell et al., 2020; Kargl et al., 2017; Horvath et al., 2024) and reflected by the positive coefficient from DM regression. Epidemiologic studies have shown that cigarette smoking leads to an increased prevalence of class-switched memory B cells in peripheral blood and memory IgG+ B cells in the lungs (Brandsma et al., 2009, 2012; Qiu et al., 2017). This is consistent with a positive relationship between the abundance of B cells and smoking in the DM regression.

## 6. Conclusion

In this study, we introduce and validate a regularized Bayesian Dirichlet-multinomial regression model for the integrative analysis of clinical information from patients and their scRNA-seq data. Our findings underscore the model’s robustness in elucidating complex interactions between cell type abundance derived from scRNA-seq data and various clinical covariates across multiple disease states. Significantly, our model successfully identified key associations between specific cell types and clinical variables in the contexts of pulmonary fibrosis, COVID-19, and lung cancer. These relationships, some of which were not previously documented, highlight the potential of single-cell technologies coupled with advanced computational methods to deepen our understanding of disease mechanisms. For instance, the model’s ability to link specific epithelial and immune cell dynamics with clinical outcomes in pulmonary fibrosis and COVID-19 provides new insights into their roles in disease progression and immune response, respectively. The hierarchical tree structure utilized in our analysis further refines the understanding of cellular interactions by mapping out how groups of cells are influenced by clinical features. This approach not only facilitates a more granular analysis but also highlights potential cellular targets for therapeutic intervention. While the model demonstrates significant promise, it also presents challenges, primarily in computational demand and data requirements. Future work will focus on optimizing these computational aspects and expanding the model’s applicability to incorporate dynamic analyses for chronic conditions, where understanding temporal changes in cell-type abundance could be particularly informative. Overall, our study advances the field of integrative analysis by providing a powerful tool for uncovering the nuanced relationships between cellular composition and clinical characteristics, thereby aiding in the development of targeted therapies and improving our understanding of complex diseases.

## Data availability

The code and data of this study are accessible through the GitHub repository at https://github.com/yg2485/Bayesian-DM-Regression.

